# Feed restriction in mid-lactation dairy cows. I: Effects on milk production and energy metabolism-related blood metabolites

**DOI:** 10.1101/2020.06.12.140996

**Authors:** I. Ansia, Y. Ohta, T. Fujieda, J. K. Drackley

## Abstract

The aim of the study was to describe the metabolic responses on energy metabolism to a period of negative nutrient balance induced by feed restriction (FR). Seven multiparous Holstein cows (93 ± 15 days in milk) were randomly assigned to 7 treatments in a 7 × 4 Youden square design. Daily intake was restricted to provide 60% of energy requirements during 5 d except for one treatment with ad libitum (AL) feeding. While 5 out of 7 experimental treatments involved abomasal supplementation of amino acids or glucose, in this article we evaluated only the effects of a negative nutrient balance by comparing both control treatments (AL and FR). Data of 2 cows within the AL group were removed due to sickness and therefore it had n = 2. Milk and energy corrected milk yield were reduced by FR. Yields of milk protein and lactose were lower during FR than during AL but the yield of milk fat only had a tendency (*P* > 0.06) to be lower with FR. Milk protein concentration was lower with FR than with AL but concentration of milk lactose and fat were not different between diets. The FR induced a decrease in plasma insulin and glucose concentrations, with quadratic decreasing trends both reaching nadirs on d 3. Simultaneously, non-esterified fatty acids (NEFA) concentration was greater and increased quadratically, peaking at d 3 during FR. There were no differences in daily β-hydroxybutyrate concentration, but it increased linearly until d 4 with FR. Comparison of the variation in concentration after feeding of insulin, NEFA and glucose could indicate a likely increased insulin sensitivity for peripheral NEFA uptake and a resistance for glucose uptake. This mechanism would contribute to decrease NEFA in circulation and sparing of glucose for lactose synthesis, respectively. Metabolic adaptations to a short-term reduction in dry matter intake include lipid mobilization, as well as modulation of peripheral tissue endocrine sensitivity in order to maintain yield of milk components production but prioritizing milk fat and lactose over milk protein.

**Implications:** The short-term feed restriction model described in this article can serve as an alternative to study metabolic adaptations during the transition period. The response of energy metabolism observed sets the baseline to measure the effect of nutrients supplementation and identify those candidates that will improve milk production and overall health after calving.

## Introduction

Homeorhetic changes in endocrine concentrations lie behind the inherent decrease of dry matter intake (**DMI**) around parturition (Drackley, 1999). A reduction in DMI across the transition period triggers a synchronized cascade of metabolic adaptations aimed to maintain an optimal concentration of glucose in circulation and a steady energy supply at the cellular level. Mobilization of stored lipids is a major mechanism to provide energy at the cellular level, but an excessive use of stored lipids can lead to metabolic malfunctions due to hepatic lipid accumulation and excessive ketone bodies formation (Bobe et al., 2004). Therefore, availability of other precursors of glucose and energy, such as amino acids (**AA**), plays a vital role during periods of negative nutrient balance (**NNB**; Arriola Apelo et al., 2014).

Feed restriction models do not simulate the natural NNB occurring during the transition period. However, this model has been a valid system to identify metabolic adaptations to nutrient deficiency at a molecular level, without the collateral effect of the endocrine regulation characteristic of parturition and the onset of lactation (Gross and Bruckmaier, 2015). The objective of this study was to characterize and quantify the effects of feed restriction on energy and protein metabolism (Ansia et al., unpublished results). In a companion paper we reported that during feed restriction the rapid decrease of most amino acids in plasma was paired with an increase in blood urea N with its peak on d 2 and decreasing afterwards. Fluctuation of the essayed N components in circulation suggests that protein tissue mobilization may have be the source of amino acids for catabolism to provide energy precursors (Ansia et al., unpublished results). Our hypothesis was that timing and magnitude of the variation in certain blood metabolites could highlight the most important metabolic pathways and accentuate requirements for specific AA in maintenance of milk production and basal metabolism when DMI is deficient. Our analysis sets the baseline to evaluate the effects of specific nutrient supplementation within this model.

## Materials and methods

### Animals and diets

Seven rumen-cannulated multiparous Holstein cows past peak lactation (93 ± 15 d in milk; BW = 674 ± 43 kg) were used in a 7 × 4 Youden square design with 4 experimental periods of 10 d. Seven treatments were applied to evaluate the effects of 5 different abomasal infusions of macronutrients on energy and protein metabolism during a 5-d feed restriction period. Only the results from the comparison of the 2 treatments used as a positive (ad libitum; **AL**) and negative (feed restriction; **FR**) controls are discussed in this article and will be referred to as diets. Cows during AL were fed to ensure daily orts of >15 % of the feed amount offered, and FR cows were fed to meet 60% of their net energy requirements for lactation (NE_L_) at the beginning of each period. During d 1 to d 5 of each period, cows received the corresponding diet (FR or AL) plus a daily 4-h abomasal infusion of distilled water (225 mL/h) beginning when the feed was delivered (09.00). Infusions were administered with a rotary peristaltic pump (Sentinel Enteral Feeding Pump, Alcor Scientific Inc., Smithfield, RI). During the following 5 d (d 6 to 10) all 7 cows were fed for ad libitum DMI to obtain 15% of refusals as fed, as a recovery and wash-out period.

All cows were housed in tie stalls and were milked twice daily at 0600 and 1700 in a parallel parlor during the duration of the trial. Milk was sampled and milk production was recorded at each milking. One aliquot of each daily milking sample was preserved (800 Broad Spectrum Microtabs II; D&F Control Systems, Inc., San Ramon, CA), refrigerated, and then analyzed (Dairy Lab Services, Dubuque, IA) for contents of fat, protein, milk urea nitrogen (**MUN**), lactose, and total solids. Another aliquot was frozen immediately at −20°C. Cows were adapted to the tie stalls and feed for 5 d before beginning the first treatment. Body weight was measured on d 1 of each period before feeding and also after the 5-d treatment on d 6 before feeding.

The diet, fed as a total mixed ration, was formulated according to the NRC (2001) requirements for a lactation diet (Tables 1). The ration was mixed daily and offered at 09.00. The restricted diet (60% of the NE_L_ requirements) was calculated according to the BW, daily milk production and composition, number of lactation, and days in milk. The same amount (DM basis) was offered during the following 5 d. Although not intended, the restricted diet provided ca. 60% of the essential AA (EAA) required as well (Table 2). The DM content of the total mixed ration was estimated daily (Koster Moisture Tester, Koster Crop Tester, Inc., Strongsville, OH; Oetzel et al., 1993) before feed delivery in an attempt to offer accurately the same relative amount of DM throughout the experiment. The total mixed ration was sampled every second day and analyzed by wet chemistry (Dairy One Cooperative Inc., Ithaca, NY); forages and concentrate components were sampled and analyzed weekly.

**Table 1.** Ingredient composition of the ration

| Item | Value |
| --- | --- |
| Ingredient (g/kg DM) |  |
| Corn silage | 342 |
| Corn grain, ground | 170 |
| Soybean hulls | 73 |
| Soybean meal, solvent 47.5% | 59 |
| Alfalfa silage | 55 |
| Molasses beet sugar | 51 |
| Cottonseed fuzzy | 50 |
| Corn gluten feed | 48 |
| Wheat middlings | 30 |
| Limestone | 12 |
| Wheat straw | 10 |
| Lactation premix<sup>1</sup> | 100 |

<sup>1</sup> Contained (as-fed basis) 49% Soyplus, 16.4% Energy Booster®, 15% ProvAAI Advantage, 8% NaOHCO<sub>3</sub>, 3.9% Biotin, 3.8% Ca<sub>2</sub>(PO<sub>4</sub>)<sub>2</sub>, 2% NaCl, 1.9%, Min/Vit premix (contained a minimum of 5% Mg, 7.5% K, 10% S, 150 mg/kg of Se, 30 g/kg of Zn, 2,205 kIU/kg of Vit. A, 662.5 kIU/kg of Vit. D3, 22 kIU/kg of Vit. E, 0.5% MgO)

**Table 2.** Nutrient composition of the ration and content of energy, metabolizable protein, and amino acids (AA) relative to requirements on each diet.

| Item | Value | % Req.<sup>1</sup> |  |
| --- | --- | --- | --- |
|  |  | AL | FR |
| Nutrient<sup>2</sup> (% of DM) |  |  |  |
| Crude protein | 16.7 | - | - |
| Metabolizable protein | 11.3 | 115 | 57 |
| ADF | 23.4 | - | - |
| aNDFom | 33.3 | - | - |
| NFC | 35.9 | - | - |
| Starch | 23.7 | - | - |
| Fat total | 5.1 | - | - |
| Ash | 8.3 |  |  |
| NE<sub>L</sub> (Mcal/kg) | 1.54 | 126 | 59 |
| AA<sup>3</sup> (% of Metabolizable protein) |  |  |  |
| Arg | 6.65 | 146 | 80 |
| His | 2.72 | 134 | 65 |
| Ile | 4.88 | 118 | 56 |
| Leu | 7.91 | 99 | 47 |
| Lys | 6.52 | 108 | 55 |
| Met | 2.05 | 97 | 49 |
| Phe | 5.01 | 113 | 54 |
| Thr | 4.62 | 139 | 72 |
| Trp | 1.36 | 109 | 54 |
| Val | 5.53 | 118 | 57 |

<sup>1</sup> Percentage of the requirement met with the ad libitum (AL) and feed-restriction (FR) diet according with the output of the AMTS software (AMTS, LLC). aNDFom = Neutral detergent fiber after amylase treatment and ashing.
<sup>2</sup> Average nutrient composition calculated from the analysis of total mixed ration.
<sup>3</sup>Amino acid content based on ration description output from AMTS.

### Blood sampling and analysis

Indwelling catheters were inserted into a jugular vein of each cow on the day before d 1 of each period. During d 1 to d 5, blood samples were obtained at −1, 0, 3, 4, 5 and 14 h relative to feeding each day. Catheters were flushed with sterile heparinized saline every 6 h between daily sampling periods and were removed on d 5 in each period. Energy metabolism was evaluated by measuring concentrations of non-esterified fatty acids (NEFA), β-hydroxy-butyrate (BHB), insulin, glucose, total cholesterol, triglycerides, and leptin. The acute-phase response was evaluated with total protein, albumin, globulin, serum amyloid A (SAA), and haptoglobin (HG). The total alkaline phosphatase (APT), aspartate aminotransferase (AST), gamma-glutamyl transferase (GGT), total bilirubin, and glutamate dehydrogenase (GDH) values were used as indicators of liver function. More information about the assayed variables and tests performed can be found in the supplementary file.

### Statistical analysis

Two cows were diagnosed with a bacterial endotoxemia and clinical mastitis, respectively, both within the AL group. Due to the noticeable detrimental effect observed on milk production, DMI, and other blood variables, and the influence that their data had on the normality and homogeneity of the model, we decided to remove them from all statistical analysis of performance and blood constituents.

Comparisons of all variables between treatments were made using the MIXED procedure of SAS version 9.4 (SAS Institute, 2012) according to the following model:

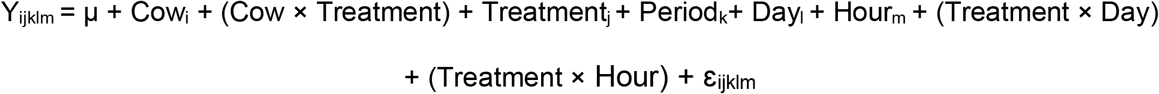

Effects of treatment, period, day, and hour were included as fixed effects in the CLASS statement while cow and the interaction cow by treatment were included as a random effects. The hour factor and its interaction with treatment were removed for variables measured only on a daily basis. The repeated fixed effect of day nested within period was used in the REPEATED statement with cow nested in treatment as the subject. Measurements of initial (d1) BW, initial milk production (kg/d), number of lactation, days in milk, and body condition score were included as covariates in the model according to their significance in explaining the variance in the model, and with their influence on the Bayesian and Akaike criteria. For each variable with equal time spacing between samples (e.g. milk, albumin, etc.), the subject was tested using 3 different covariance structures: autoregressive order 1, compound symmetry, and spatial power. For variables with unequal spacing (e.g. AA, NEFA, etc), spatial power law, Gaussian, and spherical covariance structures were tested. The covariance structure that resulted in the smallest Bayesian and corrected Akaike information criterion (AICc) was chosen (Littell et al., 2006). The degrees of freedom were estimated with the Kenward-Roger specification in the model statement.

A set of covariates was created and included to account for the carry-over effect between consecutive treatments on each variable. Covariates were retained in the model unless a 50% increase in BIC or AICc occurred or the normality of the residuals was disturbed. The CONTRAST statement was used to compare AL and FR. When the *P*-value of the interaction between treatment and day factors was lower than 0.1, the SLICE statement was applied in order to identify the specific significant interactions for each treatment for further discussion. The linear and quadratic trends were evaluated using the ESTIMATE statement from PROC MIXED, to classify the evolution of the daily average across the period.

The PROC UNIVARIATE and the INFLUENCE option within PROC MIXED were applied to each variable to check for normality of residuals and for the presence of outliers. Graph plots and *P*-values from the Shapiro-Wilk and Kolmogorov-Smirnov tests were used to evaluate both homogeneity and normality of residuals. When appropriate, variables were transformed to accomplish the above-mentioned criteria. The proper power transformation was selected according with the convenient lambda displayed by the TRANSREG procedure. Non-transformed LSMEANS and SEM were reported.

## Results

### Milk production and composition

Feed restriction reduced (*P*_Diet_ < 0.01) DMI by approximately 50 % for the FR diet in comparison with AL (Table 3). Consequently, BW difference (*P*_Diet_ = 0.03), milk yield (*P*_Diet_ < 0.01) and energy corrected milk (**ECM**: *P*_Diet_ = 0.02) were reduced during FR as well. Milk protein percentage (*P*_Diet_ = 0.04), and milk protein (*P*_Diet_ < 0.001) and lactose yields (*P*_Diet_ < 0.01) were lower with FR than AL. On the other hand, milk fat yield had a tendency (*P*_Diet_ = 0.06) to be lower with FR than AL. The total solids yield of FR was lower (*P*_Diet_ < 0.01) than AL. Milk yield and concentrations and yields of protein and lactose showed an interaction of FR with day and all manifested a decreasing linear trend across the 5-d period except total solids percentage, which tended to decrease quadratically (Table 4).

**Table 3.** Least square means of dry matter intake, energy balance, BW gain, milk production and composition of mid-lactation dairy cows during 5 days on the ad libitum (AL) and feed-restricted (FR) diets.

| Item | Diet |  | SEM<sup>1</sup> | P-values |  |  |
| --- | --- | --- | --- | --- | --- | --- |
|  | AL | FR |  | Diet<sup>2</sup> | Day | Trt x Day<sup>3</sup> |
| Dry matter intake, kg/d | 25.5 | 12.7 | 1.15 | <0.01 | <0.01 | 0.01 |
| Energy balance<sup>4</sup> (Mcal/d) | 2.5 | -12.9 | 2.4 | <0.01 | <0.01 | 0.17 |
| BW gain<sup>5</sup>, kg/d | 0.7 | -13.3 | 0.04 | 0.03 | -- | -- |
| *Yield, kg/d* |  |  |  |  |  |  |
| Milk | 39.1 | 30.9 | 1.94 | <0.01 | <0.01 | 0.05 |
| ECM<sup>6</sup> | 42.6 | 31.9 | 3.30 | 0.02 | <0.01 | 0.33 |
| Protein | 1.20 | 0.90 | 0.05 | <0.01 | <0.01 | <0.01 |
| Fat | 1.56 | 1.20 | 0.13 | 0.06 | 0.52 | 0.37 |
| Lactose | 1.82 | 1.39 | 0.09 | <0.01 | <0.01 | 0.03 |
| Total solids | 6.60 | 5.18 | 0.30 | <0.01 | <0.01 | 0.08 |
| *Milk composition* |  |  |  |  |  |  |
| Protein, % | 3.11 | 2.95 | 0.05 | 0.04 | <0.01 | 0.10 |
| Fat, % | 3.27 | 4.20 | 0.32 | 0.15 | <0.01 | 0.24 |
| Lactose, % | 4.64 | 4.60 | 0.05 | 0.78 | <0.01 | 0.34 |
| Total solids, % | 16.7 | 16.8 | 0.33 | 0.81 | 0.16 | 0.19 |
| Milk urea N, mg/kg | 136 | 140 | 12.4 | 0.82 | <0.01 | 0.82 |

<sup>1,3</sup> SEM and treatment by day interaction p-value belong to the comparison between the seven treatments.
<sup>2</sup>P-value of the orthogonal contrast between the ad libitum (AL) and feed restricted (FR) diets.
<sup>4</sup>Energy balance = NE<sub>L</sub> intake (Mcal) – (ECM × 3.14 + 0.293 × kg of BW<sup>0.75</sup>)
<sup>5</sup>BW gain between from d 1 to d 5 on each experimental period.
<sup>6</sup>ECM (kg) = (0.038 × g of fat + 0.024 × g of protein + 0.017 × g of lactose) × kg of milk/3.14.

**Table 4.** Least square means of dry matter intake, energy balance, BW gain, milk production and composition of mid-lactation dairy cows during 5 days on the ad libitum (AL) and feed-restricted (FR) diets per day.

|  |  |  |  |  |  |  |  | *P*-values |  |  |
| --- | --- | --- | --- | --- | --- | --- | --- | --- | --- | --- |
|  |  | Day |  |  |  |  |  | Trends<sup>2</sup> |  |  |
| Item | Diet | 1 | 2 | 3 | 4 | 5 | SEM | Diet x Day<sup>1</sup> | Lin. | Quad. |
| DMI, kg/d | AL | 24.5<sup>b</sup> | 25.2<sup>b</sup> | 28.1<sup>a</sup> | 27.5<sup>a</sup> | 22.0<sup>c</sup> | 1.3 | <0.01 | 0.34 | <0.01 |
|  | FR | 12.7 | 12.7 | 12.7 | 12.7 | 12.6 | 1.1 | 1.00 | 0.40 | 0.34 |
| *Yield, kg/d* |  |  |  |  |  |  |  |  |  |  |
| Milk | AL | 37.4 | 39.1 | 40.4 | 40.8 | 37.7 | 2.5 | 0.49 | 0.76 | 0.11 |
|  | FR | 35.1<sup>a</sup> | 33.1<sup>a</sup> | 30.8<sup>ab</sup> | 28.7<sup>bc</sup> | 26.7<sup>c</sup> | 1.9 | 0.01 | <0.01 | 0.99 |
| ECM | AL | 39.8 | 40.9 | 44.5 | 42.9 | 39.8 | 3.9 | 0.96 | 0.85 | 0.26 |
|  | FR | 40.1<sup>a</sup> | 33.8<sup>ab</sup> | 30.8<sup>ab</sup> | 28.9<sup>b</sup> | 26.8<sup>b</sup> | 2.9 | 0.02 | <0.01 | 0.27 |
| Protein | AL | 1.13 | 1.20 | 1.26 | 1.26 | 1.14 | 0.07 | 0.15 | 0.74 | 0.02 |
|  | FR | 1.08<sup>a</sup> | 1.00<sup>a</sup> | 0.88<sup>b</sup> | 0.80<sup>c</sup> | 0.72<sup>c</sup> | 0.05 | <0.01 | <0.01 | 0.74 |
| Fat | AL | 1.43 | 1.45 | 1.66 | 1.52 | 1.43 | 0.22 | 0.95 | 0.92 | 0.46 |
|  | FR | 1.56 | 1.23 | 1.13 | 1.08 | 1.01 | 0.15 | 0.45 | 0.02 | 0.26 |
| Lactose | AL | 1.77 | 1.84 | 1.88 | 1.88 | 1.76 | 0.12 | 0.67 | 0.94 | 0.16 |
|  | FR | 1.66<sup>a</sup> | 1.53<sup>a</sup> | 1.39<sup>b</sup> | 1.26<sup>c</sup> | 1.10<sup>d</sup> | 0.09 | <0.01 | <0.01 | 0.83 |
| Total solids | AL | 6.54 | 6.54 | 6.89 | 6.76 | 6.28 | 0.42 | 0.64 | 0.84 | 0.25 |
|  | FR | 5.86<sup>a</sup> | 5.72<sup>a</sup> | 5.21<sup>ab</sup> | 4.79<sup>bc</sup> | 4.32<sup>c</sup> | 0.34 | <0.01 | <0.01 | 0.55 |
| *Milk composition* |  |  |  |  |  |  |  |  |  |  |
| Protein, g/kg | AL | 3.06 | 3.11 | 3.16 | 3.14 | 3.07 | 0.07 | 0.53 | 0.84 | 0.10 |
|  | FR | 3.14<sup>a</sup> | 3.07<sup>a</sup> | 2.91<sup>b</sup> | 2.85<sup>bc</sup> | 2.79<sup>c</sup> | 0.06 | <0.01 | <0.01 | 0.46 |
| Fat, g/kg | AL | 3.96 | 2.97 | 3.33 | 3.00 | 3.08 | 0.60 | 0.41 | 0.39 | 0.48 |
|  | FR | 3.57 | 4.27 | 4.33 | 4.45 | 4.38 | 0.47 | 0.62 | 0.26 | 0.36 |
| Lactose, g/kg | AL | 4.65 | 4.65 | 4.59 | 4.55 | 4.61 | 0.09 | 0.60 | 0.45 | 0.49 |
|  | FR | 4.73<sup>a</sup> | 4.65<sup>ab</sup> | 4.58<sup>b</sup> | 4.45<sup>c</sup> | 4.43<sup>c</sup> | 0.07 | <0.01 | <0.01 | 0.67 |
| Total solid, g/kg | AL | 17.3 | 16.5 | 16.8 | 16.4 | 16.5 | 0.5 | 0.28 | 0.21 | 0.42 |
|  | FR | 16.8<sup>ab</sup> | 17.1<sup>a</sup> | 16.9<sup>ab</sup> | 16.9<sup>a</sup> | 16.2<sup>b</sup> | 0.4 | 0.05 | 0.17 | 0.10 |
| MUN, mg/kg | AL | 13.3 | 12.9 | 14.1 | 14.3 | 13.6 | 1.7 | 0.94 | 0.75 | 0.75 |
|  | FR | 11.8<sup>b</sup> | 15.7<sup>a</sup> | 15.1<sup>ab</sup> | 14.3<sup>ab</sup> | 13.1<sup>ab</sup> | 1.4 | 0.08 | 0.79 | 0.01 |

<sup>1</sup>P-value of the interaction of each diet (AL and FR) with day. Extracted from the SLICE statement output within the interaction of treatment with day.
<sup>2</sup>P-values of the test for linear or quadratic trends.
Means in a row with superscripts without a common letter differ, $P \leq 0.05$ .

### Energy-related metabolites and hormones in blood

Cows during FR had greater NEFA (*P* < 0.001), and lower insulin (*P* = 0.01) and glucose (*P* = 0.04) concentrations than cows during AL (Table 5). The restricted diet decreased glucose (*P*_FR x Day_ < 0.001) and insulin (*P*_FR x Day_ < 0.001) concentrations in a quadratic manner, reaching a nadir on d 3 (Table 6). The glucose-to-insulin (GIR) ratio was greater (*P*_Diet_ = 0.01) and increased (*P*_FR x Day_ < 0.01) linearly only during FR. Concentration of NEFA increased (*P*_FR x Day_ < 0.001) quadratically with its peak on d 3. Even though we did not observe an effect of diet on BHB (*P*_Diet_ = 0.33) or leptin (*P*_Diet_ = 0.52), serum BHB tended to increase linearly (*P*_FR x Day_ = 0.09) during FR. Within day, we found that hour relative to feeding had an effect on all the variables in this group with no differences between diets (Table 5). Concentrations of NEFA and glucose were lower in the samples collected during the postprandial phase (+3, +4, +5 h relative to feeding) in comparison with those pre-feeding (−1 and 0 h) and at night (+14 h), while insulin and BHB were higher (Figure 1). Changes in insulin concentration after feeding tended (*P*_Diet_ = 0.14) to be greater during FR, whereas changes in NEFA were significantly larger (*P*_Diet_ < 0.01) and changes in glucose did not vary (*P*_Diet_ = 0.46) between diets.

**Table 5.**
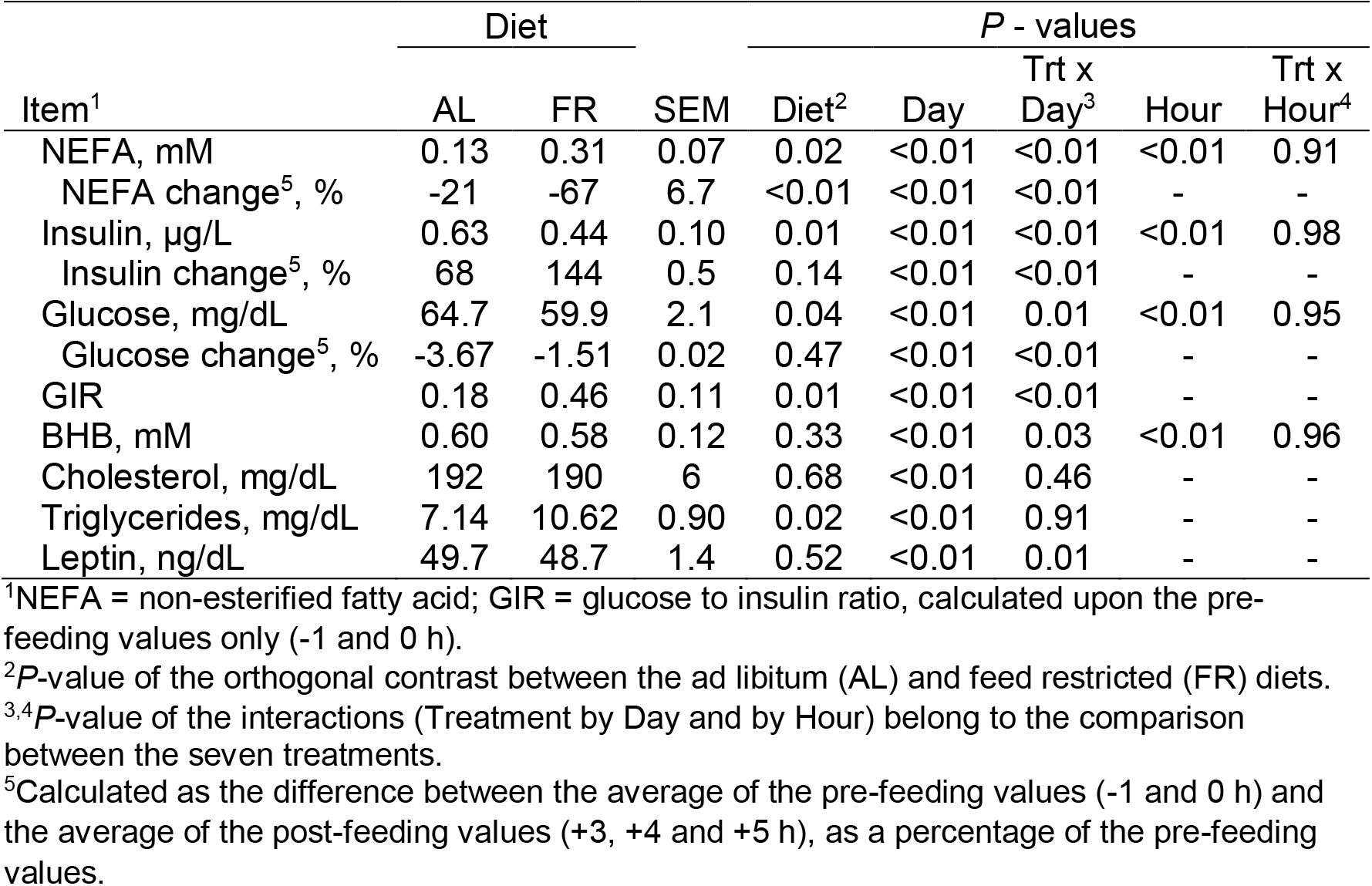
Least square means of the concentrations of blood metabolites associated with energy metabolism of mid-lactation dairy cows during 5 days on the ad libitum (AL) and feed-restricted (FR) diets.

| Item<sup>1</sup> | Diet |  |  | P - values |  |  |  |  |
| --- | --- | --- | --- | --- | --- | --- | --- | --- |
|  | AL | FR | SEM | Diet<sup>2</sup> | Day | Trt x Day<sup>3</sup> | Hour | Trt x Hour<sup>4</sup> |
| NEFA, mM | 0.13 | 0.31 | 0.07 | 0.02 | <0.01 | <0.01 | <0.01 | 0.91 |
| NEFA change<sup>5</sup>, % | -21 | -67 | 6.7 | <0.01 | <0.01 | <0.01 | - | - |
| Insulin, µg/L | 0.63 | 0.44 | 0.10 | 0.01 | <0.01 | <0.01 | <0.01 | 0.98 |
| Insulin change<sup>5</sup>, % | 68 | 144 | 0.5 | 0.14 | <0.01 | <0.01 | - | - |
| Glucose, mg/dL | 64.7 | 59.9 | 2.1 | 0.04 | <0.01 | 0.01 | <0.01 | 0.95 |
| Glucose change<sup>5</sup>, % | -3.67 | -1.51 | 0.02 | 0.47 | <0.01 | <0.01 | - | - |
| GIR | 0.18 | 0.46 | 0.11 | 0.01 | <0.01 | <0.01 | - | - |
| BHB, mM | 0.60 | 0.58 | 0.12 | 0.33 | <0.01 | 0.03 | <0.01 | 0.96 |
| Cholesterol, mg/dL | 192 | 190 | 6 | 0.68 | <0.01 | 0.46 | - | - |
| Triglycerides, mg/dL | 7.14 | 10.62 | 0.90 | 0.02 | <0.01 | 0.91 | - | - |
| Leptin, ng/dL | 49.7 | 48.7 | 1.4 | 0.52 | <0.01 | 0.01 | - | - |

<sup>1</sup>NEFA = non-esterified fatty acid; GIR = glucose to insulin ratio, calculated upon the pre-feeding values only (-1 and 0 h).
<sup>2</sup>P-value of the orthogonal contrast between the ad libitum (AL) and feed restricted (FR) diets.
<sup>3,4</sup>P-value of the interactions (Treatment by Day and by Hour) belong to the comparison between the seven treatments.
<sup>5</sup>Calculated as the difference between the average of the pre-feeding values (-1 and 0 h) and the average of the post-feeding values (+3, +4 and +5 h), as a percentage of the pre-feeding values.

**Table 6.**
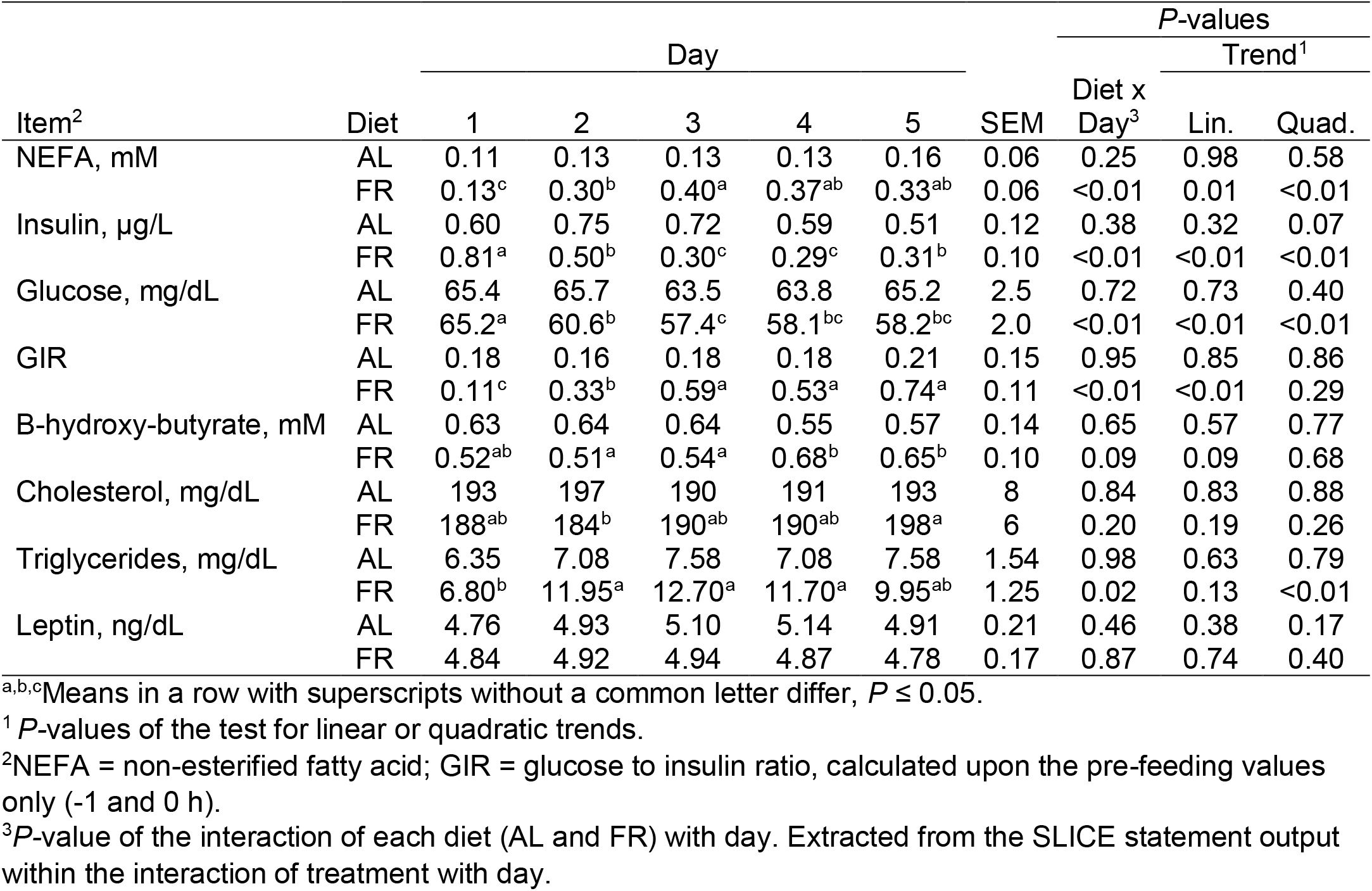
Least square means of the concentrations of blood metabolites associated with energy metabolism of mid-lactation dairy cows during 5 days on the ad libitum (AL) and feed-restricted (FR) diets per day.

**Figure 1.**
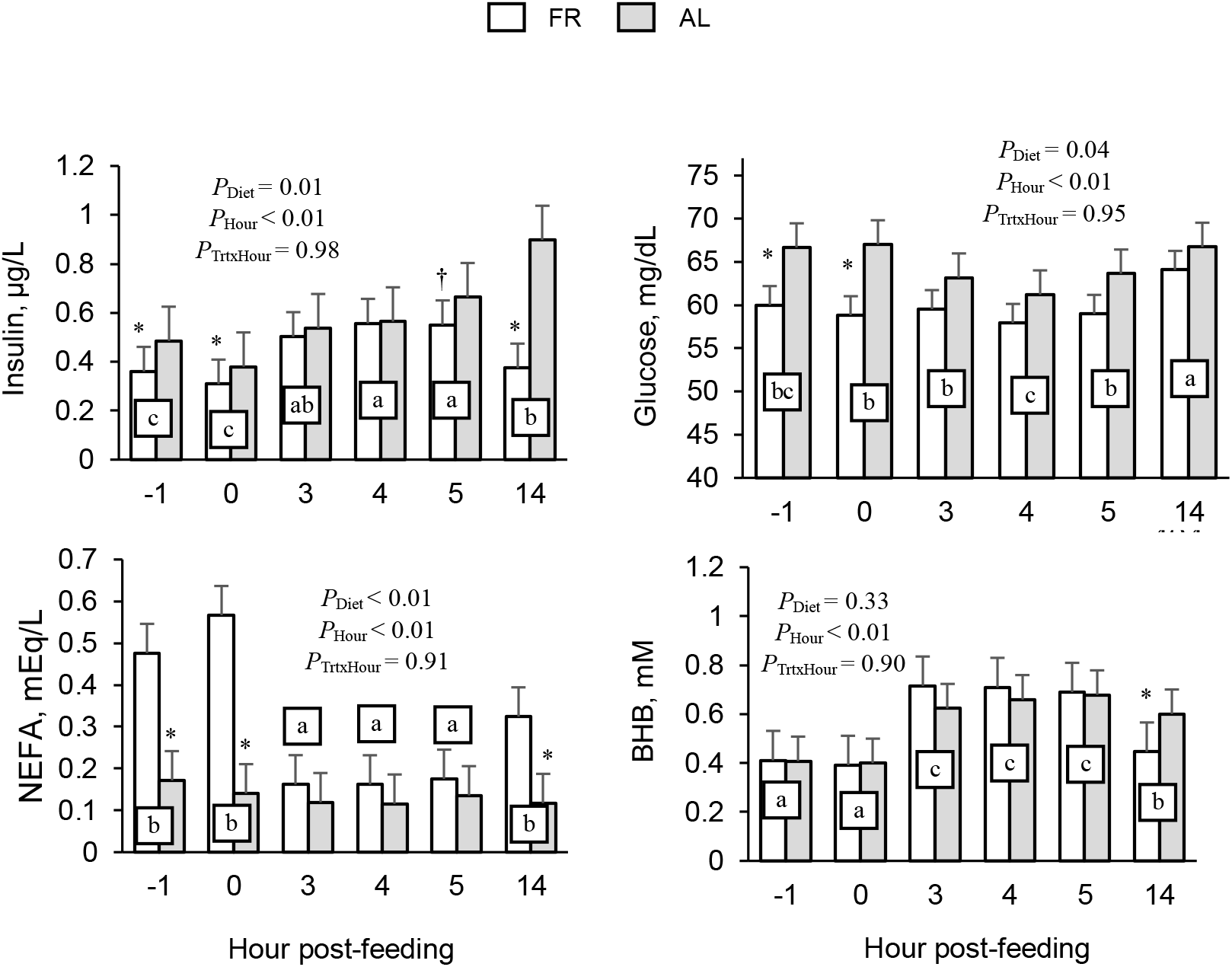
Concentrations of insulin, glucose, NEFA and BHB in plasma averaged across the 5-d period at −1, 0, +3, +4, +5 and +14 h after feeding during ad libitum (AL) and feed restricted diets (FR). Concentrations at same time points with different letter are significantly different. Differences between diets are marked at each time point with † (*P* < 0.10) and * (*P* < 0.05).

### Acute-phase response

Concentrations of albumin during FR were reduced (*P*_Diet_ = 0.01; Supplementary Table S1) following a quadratic (*P*_Quad_. = 0.03) decline across the period with its nadir on d 3 (Supplementary Table S2). Concentration of HG in AL had an interaction with day (*P*_FR x Day_ = 0.03) because its concentration on d 5 was higher than during the previous 4 d.

### Liver function variables

Among all the variables that describe liver function, we only found a profound effect of the restricted diet on bilirubin (*P*_Diet_ = 0.02) and triglycerides (*P*_Diet_ = 0.02), which were elevated in plasma during FR (Supplementary Table S1). Despite an absence of interaction between diet and day effects during FR (Supplemental Table S2), triglycerides and bilirubin had (*P*_Quad_. = 0.01) or tended to have (*P*_Quad_. = 0.12) a quadratic progression with their peak on d 3, respectively. Similarly, APT (*P*_Linear_ = 0.07) and GDH (*P*_Linear_ = 0.04) also manifested marked decreasing linear trends.

## Discussion

### Milk production and composition

Dry matter intake is the main factor that drives milk production; consequently, the reduction in milk production (−21%) and ECM (−25%) during FR was expected. The decrease in milk was smaller than the percentage in DMI reduction (−50%), which seems to indicate that metabolic adaptations were able to cope with the decreased intake. The availability of NEFA from fat mobilization (Contreras et al., 2016), the lower duodenal AA flux (Doepel et al., 2004), and the countervailing hormonal adjustments (Arriola Apelo et al., 2014) may have resulted in the alteration of milk composition during FR. We found that milk protein synthesis decreased in favor of milk fat, while lactose percentage was maintained despite the drop in blood glucose concentration. The marked decrease in protein synthesis is in agreement with Loor at el. (2007) who found that genes that regulate protein synthesis are one of the largest category downregulated during ketosis induced by feed restriction during the early postpartum. Abundance of transcripts of the major proteins in milk protein is not influenced during short periods of restricted energy supply, but is dependent on the transcription factors regulated by the endocrine system and nutrient supply (Sigl et al., 2014).

There was a lack of effect of the restricted diet on milk lactose percentage while plasma concentrations of its principal precursor glucose were reduced. This effect could imply that other anabolic mechanisms were promoting lactose synthesis (Lemosquet et al., 2009). Such mechanisms could include driving glucose towards glycolysis and lactose synthesis instead of its incorporation into milk fatty acids (Chaiyabutr et al., 1980), or by a greater insulin resistance in insulin sensitive tissue that would allow higher uptake by the mammary gland (Gross et al., 2011).

### Lipid mobilization and insulin sensitivity

The synchrony of the decline in daily concentrations of insulin and glucose, with the increase of NEFA and triglycerides in circulation indicates that d 3 was the inflection time point where metabolic homeostasis seemed to reach a stable state. Lipolysis of triglycerides in excess of re-esterification of NEFA within adipocytes is the source of NEFA in blood during periods of fat mobilization, and both processes are regulated by the antilipolytic effect of insulin (De Koster and Opsomer, 2013). These two compounds (NEFA and insulin) also reacted to feeding but in opposite directions, with no differences between diets. However, we found a difference in the magnitude of the increases in insulin (2x greater than AL), against the decreasing changes in NEFA (6x greater than AL) after feeding during the FR diet. This contrast may reflect a different muscle and adipose tissue sensitivity to insulin depending on the nutritional or energy balance status (Marett et al., 2018a), and also may provide evidence that mammary uptake of NEFA is proportional to plasma arterial concentrations (Miller et al., 1991). Similar conclusions were deduced by Piantoni et al. (2015) who found no variations of NEFA 4 h after feeding in cows in late lactation, but a marked effect in cows during their post-partum period. The different reactions between insulin and NEFA revealed that lipolytic and antilipolytic mechanisms are heavily influenced by the hourly feeding behavior (Allen et al., 2005). Furthermore, the endocrine sensitivity of tissues (Ramos-Roman et al., 2012) has more relevance than hormone concentration during periods of NNB other than the postpartum (Gross and Bruckmaier, 2015).

Liver triglycerides infiltration post-calving takes place after NEFA concentration reaches its maximum concentration (Ohgi et al., 2005). In our study, the plasma concentration of triglycerides also decreased after concentration of NEFA reached its peak in blood, suggesting that hepatic infiltration of triglycerides might have started around d 3. The NEFA recently re-esterified into triglycerides must be exported from the liver. Despite ruminant species having a low capacity for triglycerides export from liver in comparison with non-ruminants (Pullen et al., 1990), we found a weak linear increase in the concentration of cholesterol, which suggests that cows under FR still maintained a proper hepatic function as expected in mid-lactation cows (Gross et al., 2015).

Even though cows in later stages of lactation have a less intense reaction to increase ketone bodies formation (Gross and Bruckmaier, 2015), we found that daily BHB average reached its maximum at d 4, 1 day after the peak of NEFA concentration occurred. This asynchrony follows the biologic process previously mentioned wherein triglycerides infiltration and ketone bodies formation likely starts when NEFA uptake exceeded the capacity for triglycerides export and complete oxidation. In the fed state, BHB is the main ketone body synthesized by the liver and rumen epithelium. However, in the fasted state the BHB to acetoacetate ratio decreases (Heitmann et al., 1987) and acetoacetate can account for 20 to 30% of the total ketone body production in the liver (Katz and Bergman, 1969). Therefore, the magnitude of hepatic ketone body production in FR likely was underestimated.

An intense lipid mobilization would increase albumin demand to facilitate NEFA uptake into tissues (Reist et al., 2003). The quadratic decline observed for albumin points out the close relationship between NEFA and albumin as its carrier during periods of fat mobilization (Seifi et al., 2007). On the other hand, serum albumin belongs to the negative acute-phase proteins, whose production can be diminished by the effect of cytokines (Bertoni et al., 2008). Cytokines are released around calving due to diseases, stressors, or even nutritional imbalances (Drackley et al., 2005). Cytokines have a major effect in liver characterized by the induction of positive acute phase protein synthesis (e.g., haptoglobin) and the impairment of negative acute phase synthesis (e.g., albumin; Bertoni et al., 2008). Furthermore, NEFA in serum also competes with bilirubin for binding with albumin to be transported towards tissues such as the liver (Listowsky et al., 1978). Therefore, the greater concentrations of bilirubin in FR may be due to an impaired hepatic clearance due the lower availability of albumin (Reid et al., 1983). In fact, there is a positive correlation of bilirubin in serum with liver fat infiltration (West, 1990) and inflammation (Bertoni et al., 2008) in the transition period. Nevertheless, this quick response of bilirubin to lipid mobilization makes it also a good indicator of liver function during short periods of deprived feeding.

During the postprandial period, VFA from ruminal fermentation stimulate insulin release, which will drive glucose toward insulin-sensitive cells (Allen et al., 2005), thereby decreasing glucose concentration in plasma. However, the slightly greater changes in insulin concentration after feeding during FR were not accompanied by proportional decreasing changes in glucose concentration. This could indicates that, even during a short-term of NNB, there is a reactive mechanism that reduces insulin sensitivity in peripheral tissue (Marett et al., 2018b). This mechanism was also reflected by a greater and increasing GIR (calculated only in samples before feeding) during FR, which indicates that there could be a continuous state of peripheral insulin resistance (Holtenius et al., 2003). The objective of this increase in insulin resistance would be to allow a higher glucose uptake by the mammary gland when nutrients are scarce (Gross et al., 2011).

## Conclusions

The decrease in insulin concentration during FR coordinately triggered lipid and AA mobilization and oxidation as reflected by the greater NEFA and BUN concentrations in plasma and milk. Protein tissue mobilization and modulation of tissue insulin sensitivity may be important mechanisms to supply N and energy substrates for metabolic fuel, and NEFA and glucose for milk synthesis, respectively, during a short period of restricted DMI. Results from our model highlight the need for more research to better understand the role of protein tissue catabolism and endocrine regulation in shorts periods of NNB.

## Acknowledgments

The authors thank Ajinomoto Co., Inc. Tokyo, Japan for providing the funds for this experiment.

## Ethics statement

All procedures involving animals were approved by the Institutional Animal Care and Use Committee at the University of Illinois at Urbana-Champaign (protocol #15167).

