## Supplemental file for "Feed restriction in mid-lactation dairy cows. I: Effects on milk production and energy metabolism-related blood metabolites"

### Supplemental materials and methods

#### *Blood sampling and analysis*

Samples were transferred into evacuated tubes (BD Vacutainer, BD and Co., Franklin Lakes, NJ) containing clot activator for serum and $K_2EDTA$ for plasma. After blood collection, tubes for plasma were placed on ice and tubes for serum were kept at room temperature until centrifugation (~15 min). Serum and plasma were obtained by centrifugation at 2,000 × *g* for 15 min at 4°C. Aliquots of serum and plasma were frozen (−20°C) until further analysis.

Concentrations of non-esterified fatty acids (NEFA), β-hydroxy-butyrate (BHB), total protein, albumin, total alkaline phosphatase (APT), aspartate aminotransferase (AST), gamma-glutamyl transferase (GGT), total bilirubin, total cholesterol, glutamate dehydrogenase (GDH), and triglycerides were determined at the University of Illinois College of Veterinary Medicine diagnostic laboratory (Urbana, IL) using a Beckman Coulter AU680 analyzer. Total globulin was calculated as the difference between total protein and albumin. Insulin, leptin, serum amyloid A (SAA), and haptoglobin (HG) concentrations were analyzed using commercial ELISA (enzyme-linked immunosorbent assay) kits from Mercodia (catalog no. 10-1201-01), MyBioSource (catalog no. MBS703026), and Tridelta Development Ltd. (catalog no. TP 802 and TP 801), respectively.

Variables associated with the acute-phase response (total protein, albumin, HG, SAA) and liver function (APT, AST, GGT, bilirubin, GDH, cholesterol, and triglycerides) plus leptin were only analyzed in one sample (09.00) per day. The incremental changes of NEFA, glucose, and insulin concentrations after feeding were calculated as the difference between the basal concentration (average at -1 and 0 h) and the concentration during the postprandial phase (average at +3, +4 and +5 h), expressed as a percentage of the basal concentration.

### Supplemental tables

**Supplementary Table S1.** Concentrations of blood metabolites associated with acute-phase response and liver function during the ad libitum (AL) and feed-restricted (FR) diets.

| | Diet | | | *P*-values | | |
|---|---|---|---|---|---|---|
| *Item* | AL | FR | SEM | Diet[1] | Day | Trt x Day[2] |
| *Acute-Phase Response* | | | | | | |
| Total protein, g/dL | 7.83 | 7.70 | 0.12 | 0.40 | <0.01 | 0.44 |
| Albumin, g/dL | 3.29 | 3.12 | 0.05 | 0.01 | 0.49 | 0.05 |
| Globulin, g/dL | 4.57 | 4.50 | 0.09 | 0.54 | <0.01 | 0.45 |
| Haptoglobin, mg/ml | 0.80 | 0.74 | 0.16 | 0.48 | 0.23 | 0.12 |
| Serum amyloid A, ng/ml | 44.8 | 74.1 | 39.0 | 0.91 | 0.44 | 0.83 |
| *Liver Function*[3] | | | | | | |
| APT, U/L | 39.4 | 36.7 | 1.8 | 0.21 | 0.01 | 0.60 |
| AST, U/L | 71.5 | 73.1 | 4.5 | 0.78 | <0.01 | 0.27 |
| GGT, U/L | 28.5 | 28.5 | 0.7 | 0.98 | <0.01 | 0.04 |
| GDH, U/L | 50.8 | 58.9 | 22.8 | 0.40 | <0.01 | 0.92 |
| Bilirubin, mg/dL | 0.17 | 0.26 | 0.04 | 0.02 | <0.01 | 0.16 |

[1]*P*-value of the orthogonal contrast between the ad libitum (AL) and feed restricted (FR) diets.

[2]*P*-value of the interactions Treatment by Day belong to the comparison between the seven treatments.

[3] APT = Alkaline phosphatase; AST = Aspartate aminotransferase; GGT = γ- glutamyl transferase; GDH = Glutamate dehydrogenase.

**Supplementary Table S2.** Concentrations of blood metabolites associated with acute-phase response and liver function during the ad libitum (AL) and feed-restricted (FR) diets per day.

| Item | Diet | Day 1 | Day 2 | Day 3 | Day 4 | Day 5 | SEM | *P*-values: Diet x Day[2] | *P*-values: Trend[1] Lin. | *P*-values: Trend[1] Quad. |
|---|---|---|---|---|---|---|---|---|---|---|
| *Acute-Phase Response* | | | | | | | | | | |
| Total protein, g/dL | AL | 7.78 | 7.98 | 7.73 | 7.63 | 7.93 | 0.18 | 0.43 | 0.94 | 0.58 |
| | FR | 7.68 | 7.6 | 7.65 | 7.48 | 7.7 | 0.14 | 0.53 | 0.86 | 0.43 |
| Albumin, g/dL | AL | 3.33 | 3.33 | 3.18 | 3.28 | 3.33 | 0.1 | 0.77 | 0.83 | 0.27 |
| | FR | 3.22 | 3.12 | 3.05 | 3.05 | 3.15 | 0.06 | 0.25 | 0.32 | 0.03 |
| Globulin, g/dL | AL | 4.43 | 4.73 | 4.53 | 4.43 | 4.68 | 0.17 | 0.13 | 0.61 | 1 |
| | FR | 4.4 | 4.6 | 4.58 | 4.4 | 4.53 | 0.09 | 0.18 | 0.86 | 0.32 |
| Haptoglobin, mg/ml | AL | 0.64 | 0.69[a] | 0.67[a] | 0.77[a] | 1.20 | 0.23 | 0.03 | 0.11 | 0.2 |
| | FR | 0.84 | 0.86 | 0.74 | 0.7 | 0.58 | 0.18 | 0.53 | 0.24 | 0.74 |
| Serum amyloid A, ng/ml | AL | 27.1 | 43.3 | 45.2 | 47.4 | 61 | 58.5 | 0.96 | 0.7 | 0.98 |
| | FR | 78.1 | 83.8 | 72.3 | 66.1 | 70.1 | 46.4 | 0.88 | 0.82 | 0.99 |
| *Liver Function*[3] | | | | | | | | | | |
| APT, U/L | AL | 39.4 | 39.7 | 38.7 | 38.7 | 40.2 | 2.3 | 0.94 | 0.93 | 0.61 |
| | FR | 37.8 | 39.1 | 37.1 | 35.1 | 34.6 | 1.9 | 0.16 | 0.07 | 0.48 |
| AAT, U/L | AL | 74.6 | 74.2 | 70.2 | 69.7 | 69.2 | 6.1 | 0.91 | 0.48 | 0.85 |
| | FR | 74.3 | 75.8 | 76.5 | 71 | 67.8 | 4.7 | 0.47 | 0.27 | 0.26 |
| GGT, U/L | AL | 29.1 | 30.1[a] | 28.6[a] | 26.1[b] | 28.6 | 1.1 | 0.06 | 0.8 | 0.8 |
| | FR | 28.2 | 28.6 | 28.9 | 27.5 | 29.1 | 1.1 | 0.42 | 0.19 | 0.61 |
| GDH, U/L | AL | 58.4 | 58.1 | 55.4 | 56 | 51.8 | 24.6 | 0.93 | 0.85 | 0.94 |
| | FR | 87.7 | 83.5 | 74.8 | 56.6 | 37.8 | 18.8 | 0.41 | 0.04 | 0.42 |
| Bilirubin, mg/dL | AL | 0.2 | 0.18 | 0.13 | 0.18 | 0.18 | 0.06 | 0.32 | 0.88 | 0.5 |
| | FR | 0.17 | 0.26[a] | 0.31[a] | 0.26[a] | 0.29 | 0.05 | 0.03 | 0.15 | 0.12 |

[a,b,c]Means in a row with superscripts without a common letter differ, $P \leq 0.05$.

[1] *P*-values of the test for linear or quadratic trends.

[2]*P*-value of the interaction of each diet (AL and FR) with day. Extracted from the SLICE statement output within the interaction of treatment with day.

[3] APT = Alkaline phosphatase; AST = Aspartate aminotransferase; GGT = γ- glutamyl transferase; GDH = Glutamate dehydrogenase.
